# Midgut bacterial microbiota of 12 fish species from a marine protected area in the Aegean Sea (Greece)

**DOI:** 10.1101/2022.07.20.500800

**Authors:** Konstantinos Kormas, Eleni Nikouli, Vasiliki Kousteni, Dimitrios Damalas

## Abstract

Fish microbiome science is progressing fast, but it is biased toward farmed or laboratory fish species against natural fish populations, which remain considerably underinvestigated. We analysed the midgut bacterial microbiota of 45 specimens of 12 fish species collected from the Gyaros Island marine protected area (Aegean Sea, Greece). The species belong to seven taxonomic families and are either herbivores or omnivores. Mucosa midgut bacterial diversity was assessed by amplicon metabarcoding of the 16S rRNA V3–V4 gene region. A total of 854 operational taxonomic units were identified. In each fish species, between 2 and 18 OTUs dominated with cumulative relative abundance ≥70%. Most of the dominating bacterial taxa have been reported to occur both in wild and farmed fish populations. The midgut bacterial communities were different among the 12 fish, except for *Pagrus pagrus* and *Pagellus erythrinus*, which belong to the Sparidae family. No differentiation of the midgut bacterial microbiota was found based on feeding habits, i.e., omnivorous vs. carnivorous. Comparing wild and farmed *P. pagrus* midgut bacterial microbiota revealed considerable variation between them. Our results expand the gut microbiota of wild fish and support the host species effect as the more likely factor shaping intestinal bacterial microbiota.

## INTRODUCTION

Research on fish microbiomes is currently transitioning from a rather observational, i.e. structural diversity, to a more interventionist phase, i.e., looking for ways to manipulate specific microbiomes of animals of human interest to benefit the host and the environment [1]. Such manipulations are extremely restricted, if not impossible, for animals living in the wild. In addition, farmed animals experience a major deviation in their growth and environmental conditions compared with their natural counterparts, and this is expected to affect their microbiomes [2].

The benefits of knowing animal-associated microbiota stretch further than just enhancing our knowledge of microbial diversity and host-microbe interactions. It can also be considered a novel contribution to conservation practices, as host-associated microbiomes are now considered to be good indicators or even biosensors of environmental health or disturbance [3]. As microbial communities are highly responsive to environmental changes, an animal microbiota presents features which are indicative of environmental disturbance [4]. Regarding intestinal microbiota, dysbiosis ––an imbalanced microbiome with no or little beneficial traits for its host––can be related to disease or decreased well-being of the animals. In various environments, it is known that the risk of animal disease increases in degraded or disturbed environments [5]. As microbiota analysis is fast becoming more precise and less costly, it can be considered as a rather proactive, as opposed to reactive, conservation-assisting practice, which is more appropriate for the stable, long-term monitoring of ecosystem health [e.g. [6, 7]. Recently, microbiome science has been credited with having predictive potency for evolutionary processes of macroorganisms [8].

To date, most fish microbiome research is focused on farmed species [9] or on a very small number of laboratory animals, which prevents us from knowing even approximate upper limits of natural microbiome variation in wild fish; consequently, there is an imperative need for the microbial profiling of natural populations [10–13], especially the core microbiota and microbiome [14]. The core microbiome can provide beneficial adaptations to its hosts [14–16], while interindividual microbiome variability might act as a selective pressure mechanism for host adaptation, fitness, and evolution [17].

Enhancing the knowledge of wild animals’ gut microbiota not only adds to better understanding of host-microbe interactions but it may also reveal novel biotechnological potential [18]. In this paper, we present a comparative analysis of the midgut – the part of the gut where most of the microbially-mediated nutritional process take place-bacterial community structure of 12 marine fish species from the Gyaros Island marine protected area (MPA), Aegean Sea, Greece, to identify the dominant and core microbiota members and depict similarities between midgut microbiota structure of closely related fish species. The gut microbiota of eight of the investigated fish species are reported for the first time while for one them only limited data exist from farmed specimens.

## METHODS

### Study area

Sampling took place in the marine waters around Gyaros Island (Figure 1, Table S1), central Aegean Sea, Greece, which has been an MPA since 2019 [19]. According to the Greek Ministerial Decree 389/4.7.2019, spatiotemporal access for small-scale fishing is allowed and specific exploitation activities are permitted and regulated on the island. Gyaros, also locally known as Gioura, is an unpopulated island of 23 km^2^ in the northern Cyclades complex 9 nautical miles (nmi) from the closest island, Syros. Historically, Gyaros has served as a place of exile during the Roman era and the 20th century, and after World War II until 1974, it was a concentration camp for displaced political prisoners. Afterwards, it was converted to a firing range for the Greek Navy. As a result, access to other human activity was limited or restricted, and Gyaros has been under this ‘protected’ status for more than five decades. In 2011, Gyaros and the surrounding marine area of 3 nmi was included in the list of European Natura 2000 Network sites.

**Figure 1.**
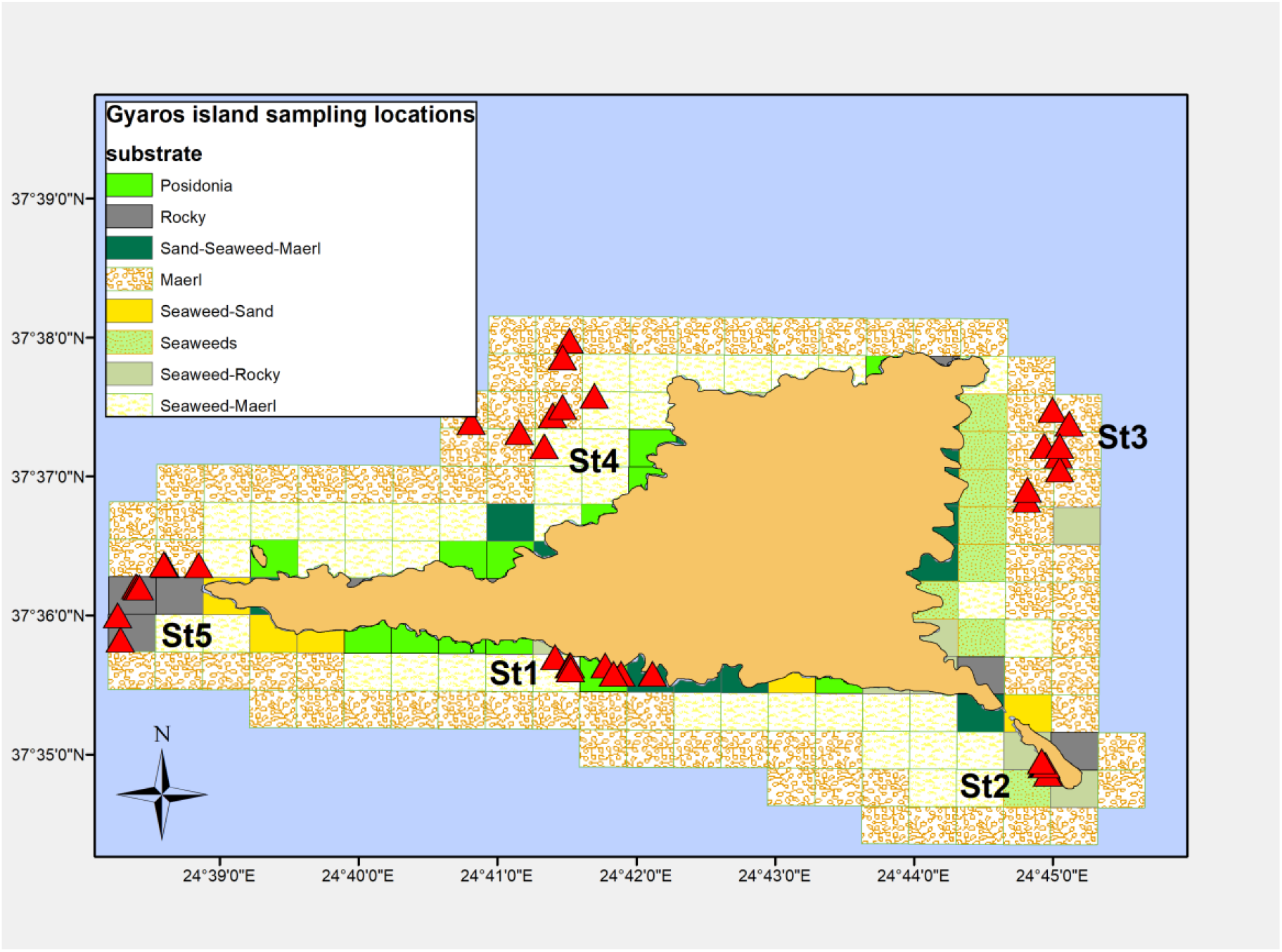
Map showing the sampling stations (St#) off Gyaros Island, Greece, in the eastern Mediterranean Sea. Red triangles indicate sampling locations.

### Fish sampling procedure

Sampling was carried out on board chartered commercial fishing vessels in five fixed stations around Gyaros Island. The sites were selected by applying NOAA’s Sampling Design Tool (https://coastalscience.noaa.gov/project/sampling-design-tool-arcgis/) and taking into account the depth strata and bottom substrate types (Figure 1, Table S1). Sampling was conducted as an experimental fishing survey from July 2018 to June 2019. Fishing gear consisted of static trammel nets with a mesh size greater than 32 mm, length from 500 to 1000 m and a height of around 2 m. The nets were cast in the late afternoon and retrieved early the next morning at sunrise. Depths over which fishing took place varied between 17 and 98 m.

Immediately after the trammel nets were hauled back to the vessels, entangled fish were extracted from the netting and identified down to the lowest possible taxonomic level. In order to avoid erroneous misidentification on board the fishing vessel, all specimens were transferred to the Hellenic Centre for Marine Research (HCMR premises), where they were thoroughly examined and identified at the species level by using specialised fish taxonomic keys based on morphometric features [20–22]. The feeding habit of each fish was determined according to Richards & Dove [23], Cortés [24], Papoutsoglou & Lyndon [25], Stergiou & Karpouzi [26], Karachle & Stergiou [27, 28] and Kousteni et al. [29]. Fish dissection took place in aseptic conditions on board fishing vessels, by using sterile gloves and scissors and scalpel were sterilized with 70% ethanol before each dissection. The entire digestive tract was excised from 45 specimens and placed in individual sterile Falcon tubes (50 mL). Both the gut samples and the remaining bodies were labelled properly, stored in DNA/RNA Shield (Zymo Research, Irvine, CA, USA) and stored in the vessels’ freezers (−20°C). After landing, the samples were transferred to laboratory facilities, where gut samples were stored at −80°C and whole bodies were frozen at −20 °C for further processing. The 45 fish specimens used for this study were clustered in seven families and 12 species. Details regarding standard biometric measurements of the specimens with corresponding metadata can be found in the supplementary material (Table S2).

### Bacterial microbiota analysis

After the gut samples were thawed on ice, the midgut section was separated and its digesta was mechanically squeezed out using forceps and excluded from further analysis. Midgut tissue samples (ca. 0.25 g) were rinsed three times with sterile, particle-free sea water. Total DNA was isolated using the QIAGEN QIAamp DNA Mini Kit (Qiagen, Hilden, Germany), following the manufacturer’s protocol “DNA Purification from Tissues”. From the extracted total DNA, bacterial DNA was amplified with the primer pair S-D-Bact-0341-b-S-17 and S-D-Bact-115 0785-a-A-21 [30] targeting the V3– V4 regions of the 16S rRNA gene. The amplified sequences were sequenced on a MiSeq Illumina instrument (2×300 bp) at the MRDNA Ltd. (Shallowater, TX, USA) sequencing facilities.

### Data analysis

Raw DNA sequences can be found in the Sequence Read Archive (https://www.ncbi.nlm.nih.gov/sra/) under BioProject PRJNA835803. The raw 16S rRNA sequencing data were processed using the MOTHUR standard operating procedure (v.1.45.3) [31, 32] and the operational taxonomic units (OTUs) at 97% cut-off similarity level, were classified with the SILVA database release 138 [33, 34]. Identification of the closest relatives of the resulting OTUs was performed using a Nucleotide Blast (http://blast.ncbi.nlm.nih.gov). Statistical analysis––including cluster analysis based on the unweighted pair group method with arithmetic mean Bray-–Curtis similarity, –permutational multivariate analysis of variance (PERMANOVA) to detect differences between the midgut bacterial microbiota of the 12 fish species regarding their OTUs composition and between the four feeding habits, and graphic illustrations were performed using Palaeontological STudies (PAST) software [35] and the vegan package [36] in R Studio platform Version 1.1.419 [37] with 3.4.3 R version.

In our study, most of the investigated fish individuals were of similar age (Table S2). However, due to the low number of individuals per fish species, we pooled individual samples from the same fish species as they had 9% to 25 % overlapping OTUs (Figure S1), to provide a more inclusive view of the core midgut bacterial diversity for each of the 12 fish species. In this study, we define as core microbiota, the shared most abundant bacterial OTUs between all 12 investigated fish species (cumulative relative abundance ≥70% after pooling together all individual samples per fish species).

## RESULTS

From the amplicon sequencing of the 16S rRNA gene V3–V4 region, a total of 3,475,413 sequences were obtained after quality filtering and chimera removal from the 45 gut samples, ranging between 160,531 and 23,517 sequences/sample. These sequences were assigned to 854 unique OTUs at a cut-off level of 97%. Sequencing data were rarefied to be equal to the smallest number (23,517) of sequences per sample.

Fish midgut bacteria at the phylum level were characterized by the predominance of Proteobacteria (61.1% relative abundance). Of the rest of the 24 detected phyla in the whole dataset, only seven (Firmicutes, Bacteroidota, Actinobacteriota, Patescibacteria, Fusobacteriota, Planctomycetota, and Dependentiae) were found to occur with ≥1% relative abundance in at least one of the 12 species. Regarding the taxonomic differences in the midgut bacteria, OTUs comprising ≥70% of the relative abundance in each fish species were assigned to 31 and 2 bacterial families and orders, respectively (Figure 2). Even the dominant OTUs were assigned to different and variable taxonomic families, except for *P. pagrus* and *P. erythrinus*. Finally, the taxonomic affiliation of these OTUs at the genus level -or higher known- is shown in Figure S2. Rank abundance curves (RACs) were also dissimilar (Figure 3).

**Figure 2.**
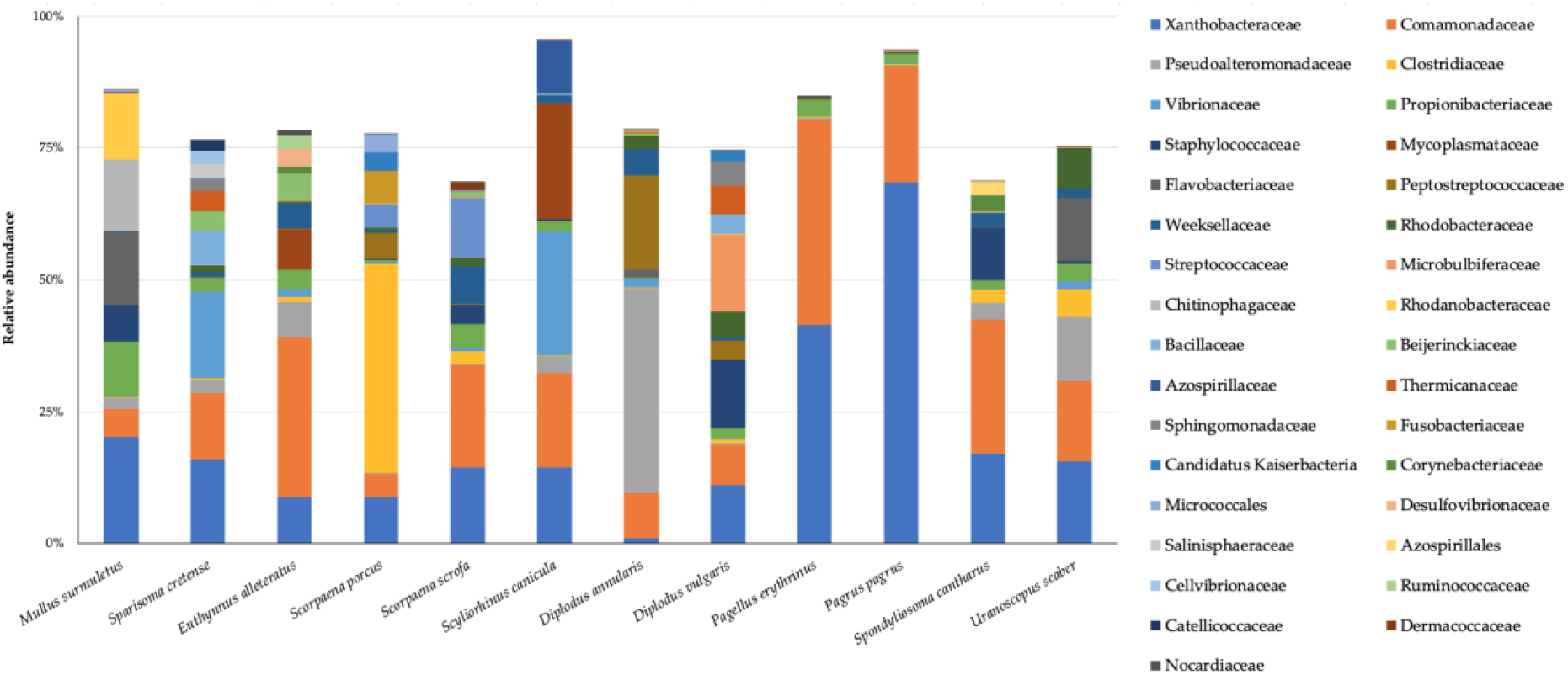
Taxonomic composition (31 families and two orders) of the most dominant (≥70% cumulative relative abundance) bacterial operational taxonomic units of the 12 fish species. Taxa in the legend are shown in decreasing total abundance in the whole dataset.

**Figure 3.**
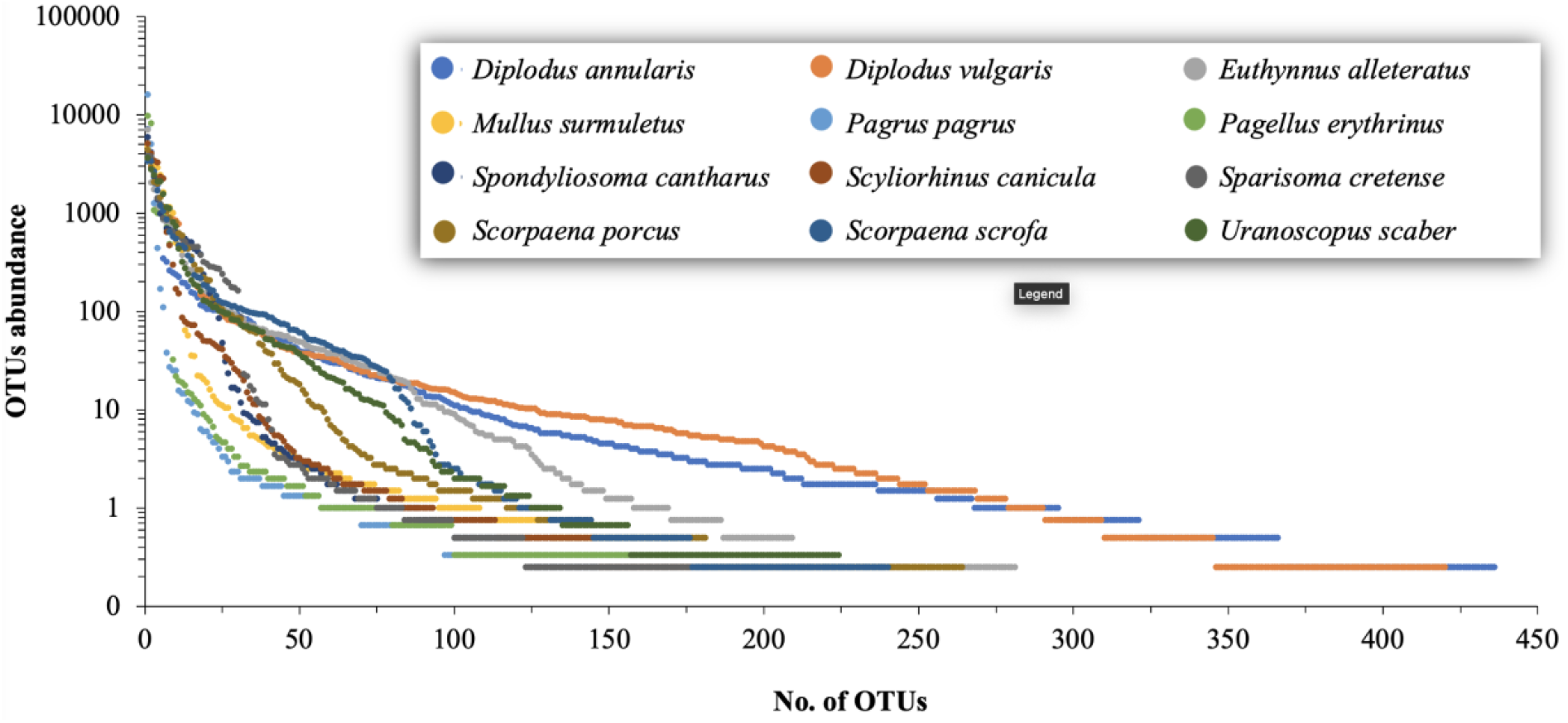
Rank abundance curves of the midgut bacterial operational taxonomic units (OTUs) abundance in 12 marine fish species from the Aegean Sea.

As core microbiota are considered functional when they are abundant [14], we present the core microbiota of the most dominant bacterial OTUs across the 12 fish species, comprising ≥70% of the cumulative relative abundance, which consists of 28 bacterial OTUs corresponding to 23 known genera and two unclassified genera (Table S3). RACs (Figure 3) showed that only a small number of dominant OTUs were found in the midgut bacterial communities, while a much larger number of OTUs were present in low and very low relative abundance. In each species, between 2 and 18 OTUs were the most abundant OTUs (cumulative relative dominance ≥70%) (Table 1). The highest OTUs richness was found in *Diplodus annularis* and the Simpson 1-D diversity index ranged between 0.25 ± 0.26 (*P*. *pagrus*) and 0.83 ± 0.12 (*Scorpaena scrofa*) (Table 1). PERMANOVA showed that the OTUs richness was different between the 12 fish species (F = 6.065, p = 0.0005). Statistically significant differences were found between *P. pagrus* and eight fish species (PERMANOVA, p_range_ = 0.027 – 0.030) and between *Spondyliosoma cantharus* and five fish species (PERMANOVA p_range_ = 0.028–0.030) (Table S4). OTUs abundance was also statistically different among the 12 fish species (F = 1.642, p = 0.0001; Table S5). Regarding the feeding habit, statistically significant differences were found between the carnivores with a preference for fish and cephalopods and the carnivores with a preference for decapods and fish and the omnivore with preference for plant material (*Sparisoma cretense*) (Table 2). Statistically significant differences were also observed based on the fish digestive physiology (F=1.527, p=0.006). The types with pyloric ceca were different with each other, while the stomachless type differed with the short and the looped Z-shaped intestine (Table 3).

**Table 1.**
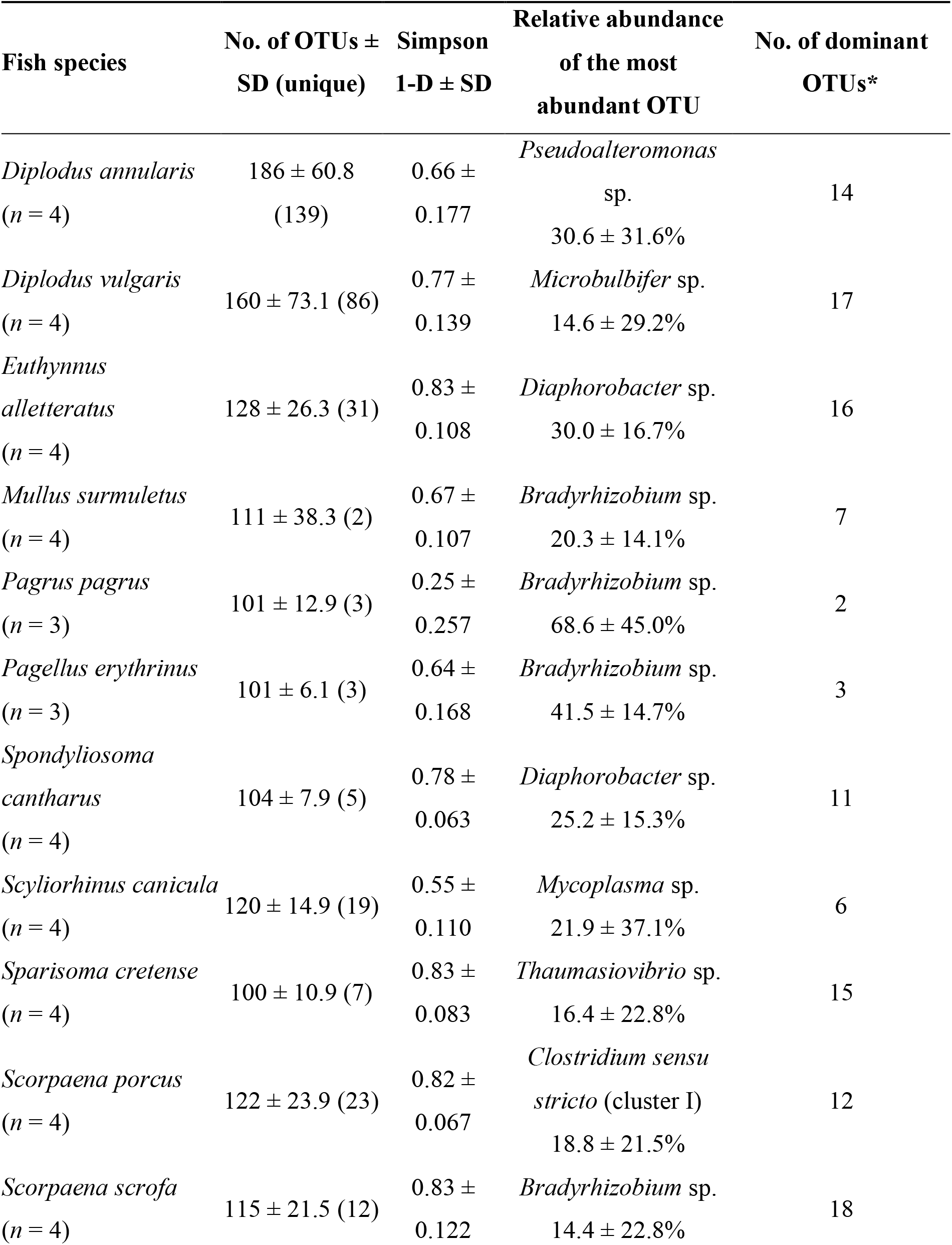
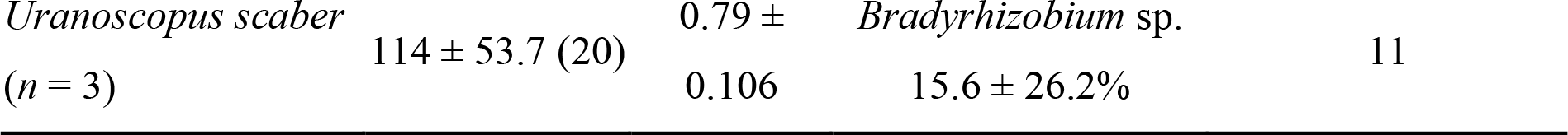
Alpha diversity data of midgut bacterial communities of 12 fish species from the Aegean Sea. SD: standard deviation. OTU: operational taxonomic unit. Unique refers to the OTUs found only in the respective fish.

**Table 2.** PERMANOVA of the midgut bacterial OTUs abundances in the midgut based on the feeding habits of 12 fish species from the Aegean Sea. Greece. Star indicates p < 0.05. OV: omnivore with a preference for plants; OA: omnivore with a preference for animal material; CD: carnivore with a preference for decapods and fish; CC: carnivore with a preference for fish and cephalopods.

|  | OA | CC | CD | OV |
| --- | --- | --- | --- | --- |
|  | *D. annularis*<br>*D. vulgaris*<br>*M. surmuletus*<br>*P. erythrinus*<br>*S. cantharus* | *E. alletteratus*<br>*S. scrofa*<br>*S. canicula*<br>*U. scaber* | *P. pagrus*<br>*S. porcus* | *S. cretense* |
| OA | - | 0.122 | 0.122 | 0.191 |
| CC |  | - | 0.013* | 0.016* |
| CD |  |  | - | 0.119 |
| OV |  |  |  | - |

**Table 3.**
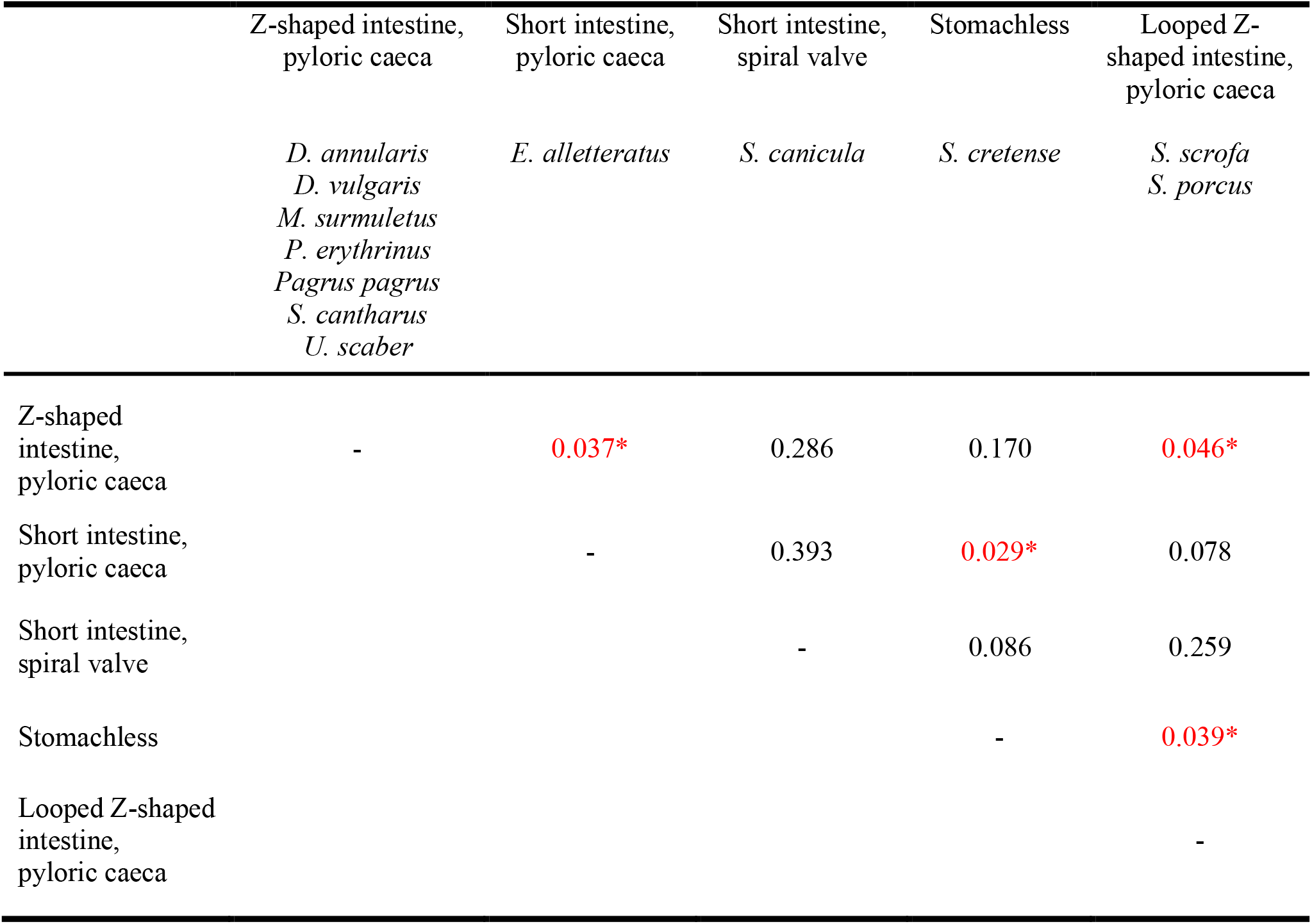
PERMANOVA of the midgut bacterial OTUs abundances in the midgut based on the digestive physiology of 12 fish species from the Aegean Sea. Greece. Star indicates p < 0.05.

|  | Z-shaped intestine,<br>pyloric caeca | Short intestine,<br>pyloric caeca | Short intestine,<br>spiral valve | Stomachless | Looped Z-shaped intestine,<br>pyloric caeca |
| --- | --- | --- | --- | --- | --- |
|  | *D. annularis*<br>*D. vulgaris*<br>*M. surmuletus*<br>*P. erythrinus*<br>*Pagrus pagrus*<br>*S. cantharus*<br>*U. scaber* | *E. alletteratus* | *S. canicula* | *S. cretense* | *S. scrofa*<br>*S. porcus* |
| Z-shaped<br>intestine,<br>pyloric caeca | - | 0.037* | 0.286 | 0.170 | 0.046* |
| Short intestine,<br>pyloric caeca |  | - | 0.393 | 0.029* | 0.078 |
| Short intestine,<br>spiral valve |  |  | - | 0.086 | 0.259 |
| Stomachless |  |  |  | - | 0.039* |
| Looped Z-shaped<br>intestine,<br>pyloric caeca |  |  |  |  | - |

There was no clear clustering pattern for species of the same family (five species in the Sparidae family and two in the Scorpaenidae family), except for *P. pagrus* and *P. erythrinus* of the Sparidae family (Figure 4). Similarly, no clustering was observed regarding feeding habits (carnivore/omnivore, see Table S6). Similar topologies were also found with UniFrac analysis (Figure S3). However, regarding the presence of shared family-level OTUs, the five Sparidae species shared 61 of 142 (42.9%) OTUs, while the two Scorpaenidae species shared 62.2% of their total OTUs (Figure 5).

**Figure 4.**
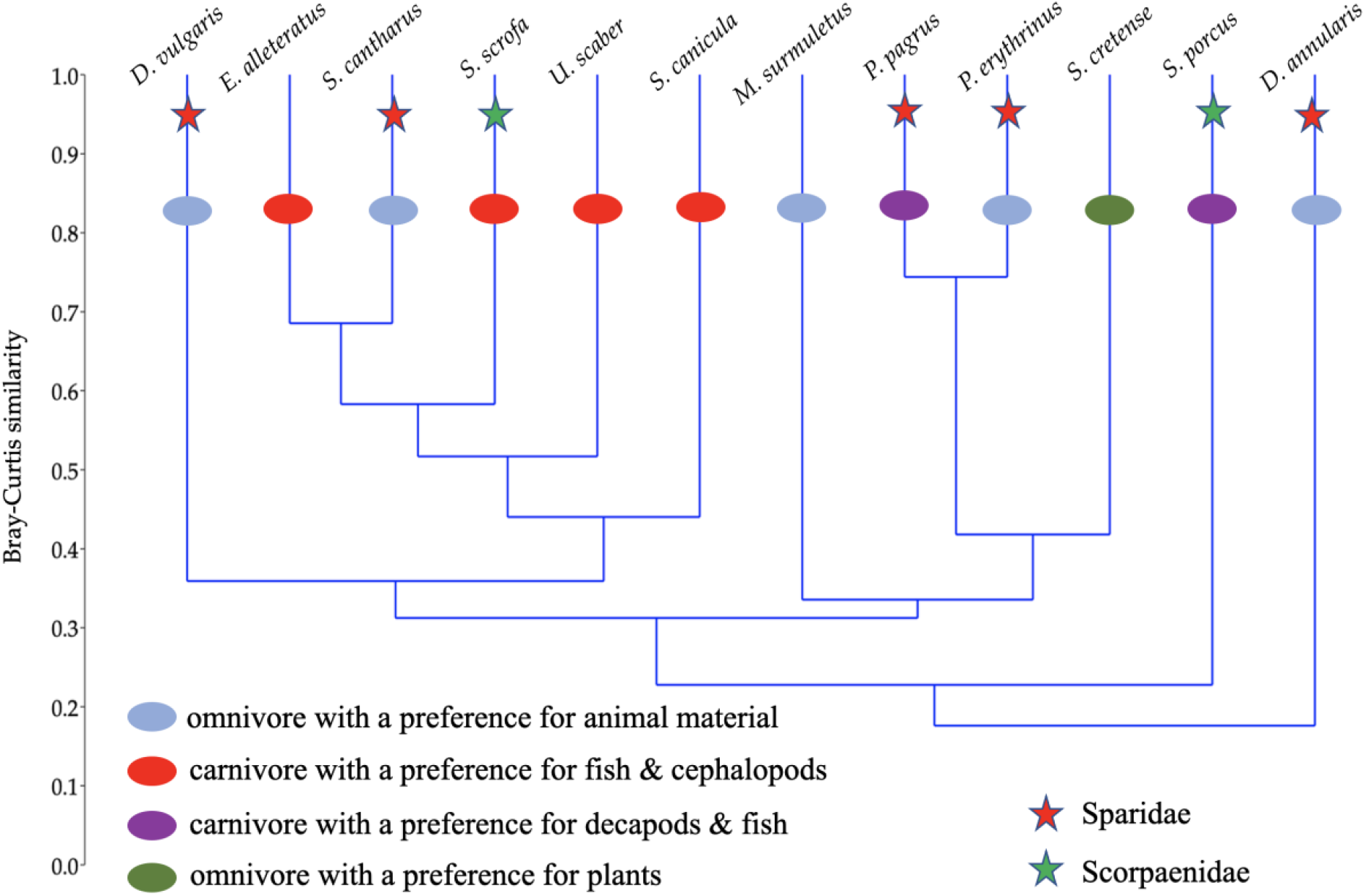
Cluster analysis of the 12 fish species’ midgut bacterial microbiota.

**Figure 5.**
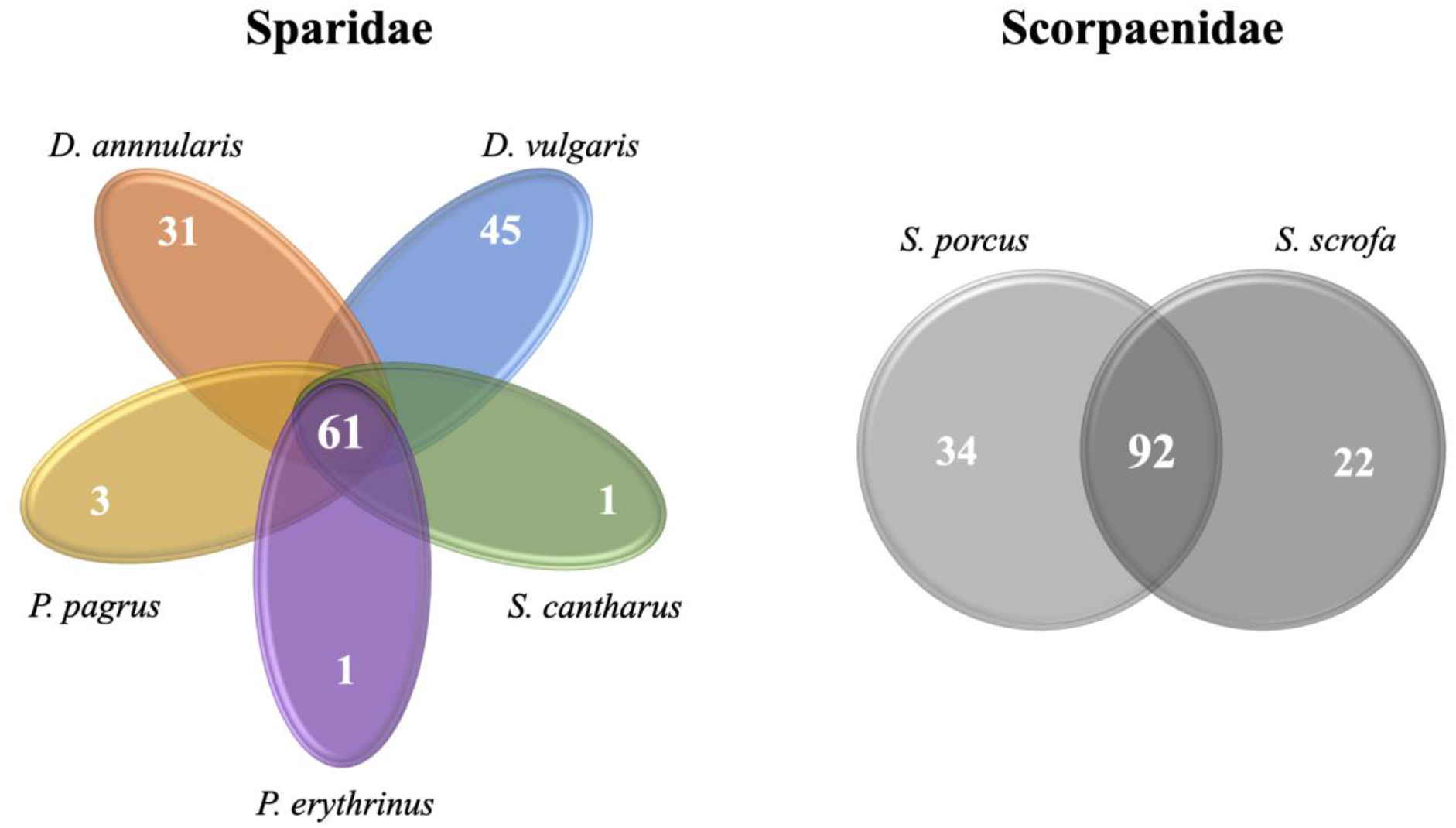
Shared (i.e. found in all samples of the fish species of each diagram) and unique (found only in the respective fish species) midgut bacterial family-level operational taxonomic units (OTUs).

## DISCUSSION

The definition of the core microbiota is based on shared microorganisms among comparable habitats [38], and for this, a large number of replicate samples was needed to overcome the effect of individual variability [39]. Indeed, such prerequisites are achievable under experimental conditions (e.g., [11, 40]) or in the case of farmed fish populations (e.g., [41–43]). However, when core microbiota of fish from natural populations are investigated by experimental fishing in the open sea, collecting adequate replicates of specimens of the same age cannot be secured, so scientists must rely on what they catch. In the present study, despite that the variation of the midgut bacterial abundance in each fish species was high -most likely due to the small number of individuals per species we managed to collect-it was considerably lower than the variation of all individuals combined for the 12 fish species (Table S7).

The core microbiota across different animal hosts is a kind of genetic link among these species since these microorganisms hold the essential core genes for different hosts [44]. Following [45], we present the core midgut bacterial microbiota of the most abundant OTUs of 12 marine fish species from the Gyaros Island marine protected area in the Aegean Sea, Greece. For eight of these species (*E. alletteratus, M. surmuletus, S. porcus, S. scrofa, S. canicula, S. cretense, S. cantharus* and *U. scaber*) there are no available data on gut microbiota and microbiomes to date, while the gut microbiota of *D. annularis, D. vulgaris, P. pagrus* and *P. erythrinus* have been recently investigated by Escalas et al. [46]. Regarding *P. pagrus*, its gut bacterial microbiota has been studied in a farmed population [43]. The core microbiota of these 12 species consisted of 23 known and two yet-unaffiliated (from the Clostridiaceae and Vibrionaceae families) genera (Table S2). The majority of these genera are known to occur in the intestinal systems of healthy wild and farmed fish species [2, 47, 48], thereby expanding their keystone and fundamental ecophysiological role across several fish species.

There were multiple differences in midgut bacteria structure among the 12 species. At first, OTU richness was highly variable, as illustrated by both the range (Table 2) and the RACs (Figure 3). The latter revealed only a small number of dominant OTUs; however, these bacteria were taxonomically different. Such a microbiota structure, i.e., a few dominant OTUs with a large “rare biosphere” [49], has been reported in other comparative studies of fish gut microbiota [10, 12]. In a comparison of wild and farmed populations of the same species, higher gut bacterial diversity in animal hosts is thought to be beneficial [50] because it sustains a functional microbiota in an environment with a fluctuating food supply [10]. The same has been proposed for other animals in the wild [18]. Due to functional redundancy of bacteria, taxonomic diversity does not necessarily reflect metabolic diversity [51]; however, the extent of functional diversity of wild fish microbiota [1] remains to be studied.

Although host genetics can be the major factor shaping fish gut microbiome [52], in closely related taxa or species with similar intestinal system development [53], host genetics may not be the most important factor distinguishing microbiome (e.g., [54]). In the present study, only two pairs of fish species had more than 70% similarity in their midgut bacterial microbiota (Figure 4) which could be attributed either to genetic relatedness or similar feeding habits. Despite this structural similarity, PERMANOVA for the first pair, *E. alletteratus* and *S. cantharus*, which belong to different taxonomic families, showed that their midgut bacterial microbiota was statistically different (PERMANOVA *p* < 0.001, F = 9.71). The second pair, *P. pagrus* and *P. erythrinus*, of the Sparidae family, are more closely related (Figure S4), as no statistically significant difference was found (PERMANOVA *p* = 0.889, F = 0.768). Both fish pairs consisted of carnivorous species with different preferences (Table S6). Our dataset included two pairs of fish species from the same genus, *D. annularis*/*D. vulgaris* and *S. porcus/S. scrofa*; however, their in-between midgut bacterial microbiota structure was very distant (Figure 4). Although host phylogeny and diet remain the two major shaping factors of animal gut microbiota [53], the above pairwise comparisons suggest that host species can shape the gut microbiota rather than diet or feeding habit of wild marine fish. In the present study, both the feeding habit and the digestive physiology of the 12 fish did not show a consistent pattern of differences with the rest of the types, i.e., there was not even a single feeding habit or digestive physiology group being different with all of the rest groups (Table 2, 3). Despite that each feeding habit and digestive physiology categories did not have similar numbers of fish species, the rather sporadic -or, at least, inexplicable with the current data-differences leave the host species effect as the more likely factor for the shaping of the midgut bacterial microbiota; additional data from more species per feeding type category are required in future studies to clarify this issue. Even in co-farmed species reared under the same environmental and dietary conditions, each fish species was found to host its own distinct gut microbiota [43]. However, gut microbiota is susceptible to manipulations at least for experimental or commercial rearing purposes [1]. In a recent study, host habitat, i.e. freshwaters vs. marine waters, was found to be the prime factor for converging microbiota structure in multiple wild fish species [12]. In the present study, all fish were caught around a small island in a marine protected area from similar habitats and substrates (Figure 1, Table S1), so habitat variability was not expected to have a large effect on the midgut bacterial microbiota.

The dominant bacterial phyla found in the current study were the expected ones based on what is known about intestinal microbiota of wild [12] and farmed marine fish [2]. *P. pagrus* and *P. erythrinus*, the only two species with the highest similarity in midgut microbiota compared to the other species, were dominated by OTUs belonging to the families of Xanthobacteriaceae and Comamonadaceae. More specifically, for both species the dominant OTU was affiliated with the genus *Bradyrhizobium*, which was also dominant in *Mullus surmuletus*, *Scorpaena scrofa* and *Uranoscopus scaber* (Table 2). *Bradyrhizobium* has been found to co-dominate with known probiotic bacteria in the midgut of the olive flounder *Paralichthys olivaceus* under experimental conditions [55]. In the gilthead seabream, *Bradyrhizobium*, along with the known probiotic *Weissella*, was favoured after a transition from a high- to low-lipid diet [56]. *Bradyrhizobium* has not been reported to be associated with fish disease, but it has been found to be sensitive to *Vibrio harveyi* infection [57]. Thus, it seems to be a fundamental, i.e., highly abundant, bacterial resident in the *P. pagrus* and *P. erythrinus* gut bacterial microbiota with potentially beneficial role to its hosts.

Most of the other dominant OTUs were affiliated with multiple bacterial genera (Table 2) that had a proven beneficial, or at least neutral, effect on their hosts. *Pseudoalteromonas* genus is often found among dominant OTUs in the healthy intestines of several wild fish species [13], so it is considered to be a beneficial microorganism for fish hosts [58, 59] even in the early life stages [60, 61]. Furthermore, we recently found that it dominates the fertilized eggs of *Seriola dumerili* in commercial hatcheries [Kormas et al. unpublished]. Fish-associated *Diaphorobacter*-related OTUs have been reported only once and found to prevail in the wild and farmed gilthead seabream midguts [62]. *Mycoplasma* dominated in the wild but not in the farmed populations of red-cusk eels [63], Atlantic salmon [64, 65], rainbow trout [66] and a few other wild-caught fish species [13]. *Clostridium* spp. are favoured in the gut of herbivorous fish [2] and are well-established probiotics in the aquaculture industry [67]. A *Microbulbifer* sp. strain was recently isolated from the intestine of the herbivorous teleost *Girella melanichthys* and was found to degrade cellulose [68], a rather useful trait for the omnivorous *D. vulgaris* midgut in the current study. *Thaumasiovibrio* is a recently described genus [69] and has not yet been reported to be associated with fish.

The captivity conditions of wild animals [e.g., [70, 71]] and even urbanization of human populations [e.g., [72]] have been shown to alter the gut microbiomes of these macroorganisms. Of the 12 fish species investigated in the present study, and according to the Hellenic Statistical Authority (https://www.statistics.gr/en/home/), only *Pagrus pagrus* is included in the list of farmed fish in Greece. The midgut microbiota of this farmed species was recently reported after being farmed sympatrically with four other fish species under the same environmental and dietary conditions, using the same sampling procedure and data processing and analysis but with lower sequencing depth (3,027 reads compared to 23,517 reads per sample in the present study) [43]. It was found that the *P. pagrus* midgut hosted only 29 bacterial OTUs, which were dominated by *Hydrogenophilus*, followed by *Pseudomonas, Stenotrophomonas, Bradyrhizobium, Hydrogenophaga, Comamonas, Propionibacterium* and *Janibacter*. All these OTUs comprised ca. 81% of the total relative abundance of all midgut bacteria. In the present study, the wild *P. pagrus* midgut had 179 OTUs with just the *Bradyrhizobium* and *Pelomonas* OTUs comprising ca. 90.1% of the relative bacterial abundance. Thus, wild and farmed *P. pagrus* midgut bacterial microbiota have a different community structure, although specimens in the current study were smaller (Table S1) from those reported in Nikouli et al. [43] (their Table S1). Such differences in the microbiota between wild and farmed populations of the same species have also been shown in targeted studies for several fish species such as gilthead seabream [62], red-cusk eel [63], fine flounder [73], Atlantic salmon [11], yellowtail amberjack [74], and large yellow croaker [75].

This study investigated the 16S rRNA gene amplicon metabarcoding of the midgut bacterial communities of 12 wild fish species collected from the Gyaros Island marine protected area in Aegean Sea, Greece. For eight of them, it is the first time their gut bacterial microbiota is reported, while for one species, only limited data exist on its farmed counterpart; it’s wild and natural populations seem to host different midgut bacterial microbiota. There was a general pattern of diverging microbiota in closely related fish species, expect for *P. pagrus* and *P. erythrinus* of the Sparidae family, which had very similar midgut microbiota. The dominant core bacterial microbiota consisted of 28 different OTUs, with most of them being related to bacterial taxa reported in other healthy wild and farmed species, some of which have a beneficial effect on their hosts. Finally, as new conservation strategies are starting to include natural microbiomes as part of ecosystem monitoring for conservation strategies, our results provide microbiota composition and structure, related mostly to the nutrition of higher trophic-level macro-organisms in a pristine and protected marine are

## ACKNOWLEDGMENTS

The sampling surveys were realized during the “Gyaros MPA fisheries knowledge survey: assessing a pristine Mediterranean biodiversity hotspot” funded by the MAVA Foundation (Grant Agreement 17114). Special thanks go to the local fishers on whose vessels we conducted all experimental fishing surveys.

## STATEMENTS & DECLARATIONS

This research is co-financed by Greece and the European Union (European Social Fund-ESF) through the operational program “Human Resources Development, Education and Lifelong Learning 2014–2020” in the context of the project “The diversity of gut microbiota of multiple Aegean Sea fish species” (MIS 5048929).

The authors have no relevant financial or non-financial interests to disclose.

Author contributions: Conceptualization, K.K.; methodology, K.K., E.N., V.K., and D.D.; formal analysis, K.K., E.N., V.K., and D.D.; data curation, K.K. and E.N.; writing— original draft preparation, K.K. and E.N.; writing—review and editing, K.K., E.N., V.K., and D.D.; project administration, K.K.; funding acquisition, K.K. and D.D. All the authors have read and agreed to the published version of the manuscript.

