## Supplementary tables and figures for "Midgut bacterial microbiota of 12 fish species from a marine protected area in the Aegean Sea (Greece)"

### Supplementary file

**Table S1.** Sampling locations off Gyaros Island. Greece. eastern Mediterranean Sea.

| Station ID | Location | Coordinates | Depth of fishing (m) | Dominant substrate type |
|---|---|---|---|---|
| St. 1 | Fyllada | 37° 35.421’ N<br>24° 42.129’ E | 18 | *Posidonia*/Rocky |
| St. 2 | Glaronissi | 37° 34.860’ N<br>24° 45.015’ E | 17 | *Posidonia*/Rocky |
| St. 3 | Fournaki | 37° 37.486’ N<br>24° 45.117’ E | 98 | Maerl |
| St. 4 | Colata | 37° 37.323’ N<br>24° 40.998’ E | 88 | Maerl |
| St. 5 | Fouis | 37° 36.367’ N<br>24° 38.612’ E | 47 | Rocky |

**Table S2**. Standard biometric measurements of the fish specimens used in the study with the corresponding metadata.

| Family | Species | Sampling information | | | | | |
|---|---|---|---|---|---|---|---|
| | | MONTH | YEAR | STATION | MATURITY STAGE | TL (mm) | $M_E$ (g) |
| Sparidae | *Diplodus annularis* | JUL. | 2018 | 2 | 5 | 163 | 68 |
| | | SEP. | 2018 | 2 | 3 | 156 | 66 |
| | | FEB. | 2019 | 2 | 3 | 171 | 29.6 |
| | | JUL. | 2018 | 2 | 3 | 152 | 52 |
| | *Diplodus vulgaris* | JUL. | 2018 | 1 | 2 | 182 | 80 |
| | | JUL. | 2018 | 1 | 2 | 167 | 53 |
| | | JUL. | 2018 | 5 | 2 | 193 | 106 |
| | | JUL. | 2018 | 2 | 2 | 202 | 121 |
| | *Pagrus pagrus* | FEB. | 2019 | 3 | NA | 312 | 409 |
| | | FEB. | 2019 | 3 | 5 | 351 | 357.4 |
| | | FEB. | 2019 | 3 | 2 | 249 | 222 |
| | *Pagellus erythrinus* | JUL. | 2018 | 4 | 4 | 306 | 339 |
| | | FEB. | 2019 | 3 | 2 | 265 | 226.3 |
| | | FEB. | 2019 | 4 | 2 | 360 | 447.9 |
| | *Spondyliosoma cantharus* | JUL. | 2018 | 5 | 2 | 225 | 193 |
| | | JUL. | 2018 | 1 | 2 | 212 | 124 |
| | | JUL. | 2018 | 5 | 2 | 221 | 157 |
| | | JUL. | 2018 | 5 | 2 | 223 | 161 |
| Scorpaenidae | *Scorpaena porcus* | SEP. | 2018 | 2 | 2 | 188 | 129 |
| | | SEP. | 2018 | 1 | 2 | 208 | 165 |
| | | SEP. | 2018 | 2 | 2 | 191 | 146 |
| | | SEP. | 2018 | 2 | 2 | 194 | 139 |
| | *Scorpaena scrofa* | JUL. | 2018 | 2 | 2 | 184 | 105 |
| | | JUL. | 2018 | 2 | 2 | 207 | 129 |
| | | JUL. | 2018 | 2 | 2 | 310 | 514 |
| | | JUL. | 2018 | 2 | 2 | 278 | 350 |
| Mullidae | *Mullus surmuletus* | FEB. | 2019 | 4 | 4 | 270 | 263.2 |
| | | FEB. | 2019 | 1 | 4 | 255 | 208 |
| | | FEB. | 2019 | 3 | 4 | 298 | 314.6 |
| | | FEB. | 2019 | 3 | 4 | 290 | 290.4 |
| Scyliorhinidae | *Scyliorhinus canicula* | JUL. | 2018 | 3 | 4 | 409 | 158 |
| | | JUL. | 2018 | 3 | 4 | 447 | 435 |
| | | JUL. | 2018 | 4 | 4 | 432 | 191 |
| | | JUL. | 2018 | 3 | 4 | 422 | 187 |
| Scaridae | *Sparisoma cretense* | JUL. | 2018 | 2 | 5 | 232 | 180 |

| | | | | | | | |
|---|---|---|---|---|---|---|---|
| | | JUL. | 2018 | 2 | 5 | 278 | 319 |
| | | JUL. | 2018 | 2 | 5 | 276 | 317 |
| | | JUL. | 2018 | 2 | 5 | 345 | 591 |
| Scombridae | *Euthynnus alletteratus* | FEB. | 2019 | 1 | 2 | 469 | 1195.4 |
| | | FEB. | 2019 | 1 | 1 | 392 | 631.8 |
| | | FEB. | 2019 | 1 | 2 | 390 | 716.6 |
| | | FEB. | 2019 | 1 | 2 | 461 | 1173.8 |
| Uranoscopidae | *Uranoscopus scaber* | SEP. | 2018 | 1 | 1 | 231 | 184 |
| | | SEP. | 2018 | 2 | 1 | 275 | 360 |
| | | FEB. | 2019 | 2 | 1 | 264 | 306.8 |

**Table S3**. Core (shaded) and most abundant (cumulative relative abundance ≥70% per sample) bacterial operational taxonomic units (OTU) in the midgut of 12 fish species from the Aegean Sea. Greece.

| *OTU-s* | *Phylum* | *Class* | *Order* | *Family* | *Genus* |
|---|---|---|---|---|---|
| OTU-001 | Proteobacteria | Alphaproteobacteria | Rhizobiales | Xanthobacteraceae | *Bradyrhizobium* |
| OTU-002 | Proteobacteria | Gammaproteobacteria | Burkholderiales | Comamonadaceae | *Diaphorobacter* |
| OTU-003 | Proteobacteria | Gammaproteobacteria | Enterobacterales | Pseudoalteromonadaceae | *Pseudoalteromonas* |
| OTU-004 | Proteobacteria | Gammaproteobacteria | Burkholderiales | Comamonadaceae | *Pelomonas* |
| OTU-005 | Firmicutes | Clostridia | Peptostreptococcales-Tissierellales | Peptostreptococcaceae | *Romboutsia* |
| OTU-006 | Firmicutes | Clostridia | Clostridiales | Clostridiaceae | *Clostridium* sensu stricto 1 |
| OTU-007 | Firmicutes | Bacilli | Staphylococcales | Staphylococcaceae | *Staphylococcus* |
| OTU-008 | Bacteroidota | Bacteroidia | Flavobacteriales | Weeksellaceae | *Cloacibacterium* |
| OTU-009 | Proteobacteria | Gammaproteobacteria | Enterobacterales | Vibrionaceae | Unclassified |
| OTU-010 | Proteobacteria | Gammaproteobacteria | Pseudomonadales | Microbulbiferaceae | *Microbulbifer* |
| OTU-011 | Actinobacteriota | Actinobacteria | Propionibacteriales | Propionibacteriaceae | *Cutibacterium* |
| OTU-012 | Firmicutes | Clostridia | Clostridiales | Clostridiaceae | *Clostridium* sensu stricto 1 |
| OTU-013 | Proteobacteria | Gammaproteobacteria | Enterobacterales | Pseudoalteromonadaceae | *Pseudoalteromonas* |
| OTU-014 | Bacteroidota | Bacteroidia | Flavobacteriales | Flavobacteriaceae | *Capnocytophaga* |
| OTU-015 | Firmicutes | Bacilli | Lactobacillales | Streptococcaceae | *Streptococcus* |
| OTU-016 | Firmicutes | Clostridia | Clostridiales | Clostridiaceae | Unclassified |
| OTU-017 | Proteobacteria | Gammaproteobacteria | Enterobacterales | Vibrionaceae | *Thaumasiovibrio* |
| OTU-018 | Firmicutes | Bacilli | Thermicanales | Thermicanaceae | *Thermicanus* |
| OTU-019 | Firmicutes | Bacilli | Mycoplasmatales | Mycoplasmataceae | *Mycoplasma* |
| OTU-020 | Proteobacteria | Alphaproteobacteria | Rhizobiales | Beijerinckiaceae | *Bosea* |

| | | | | | |
|---|---|---|---|---|---|
| OTU-021 | Firmicutes | Bacilli | Bacillales | Bacillaceae | *Aeribacillus* |
| OTU-022 | Fusobacteriota | Fusobacteriia | Fusobacteriales | Fusobacteriaceae | *Cetobacterium* |
| OTU-023 | Proteobacteria | Gammaproteobacteria | Xanthomonadales | Rhodanobacteraceae | *Luteibacter* |
| OTU-024 | Firmicutes | Bacilli | Mycoplasmatales | Mycoplasmataceae | *Mycoplasma* |
| OTU-025 | Bacteroidota | Bacteroidia | Flavobacteriales | Flavobacteriaceae | *Polaribacter* |
| OTU-026 | Proteobacteria | Alphaproteobacteria | Rhodobacterales | Rhodobacteraceae | *Ruegeria* |
| OTU-027 | Bacteroidota | Bacteroidia | Chitinophagales | Chitinophagaceae | *Puia* |
| OTU-028 | Proteobacteria | Alphaproteobacteria | Rhodobacterales | Rhodobacteraceae | *Paracoccus* |
| OTU-029 | Patescibacteria | Parcubacteria | *Candidatus* Kaiserbacteria | *Candidatus* Kaiserbacteria | Unclassified |
| OTU-030 | Firmicutes | Bacilli | Bacillales | Bacillaceae | *Anoxybacillus* |
| OTU-031 | Actinobacteriota | Actinobacteria | Micrococcales | Micrococcales *incertae sedis* | *Timonella* |
| OTU-032 | Proteobacteria | Gammaproteobacteria | Enterobacterales | Vibrionaceae | *Photobacterium* |
| OTU-033 | Actinobacteriota | Actinobacteria | Micrococcales | Bogoriellaceae | *Georgenia* |
| OTU-034 | Dependentiae | Babeliae | Babeliales | Vermiphilaceae | Unclassified |
| OTU-035 | Proteobacteria | Alphaproteobacteria | Sphingomonadales | Sphingomonadaceae | *Sphingomonas* |
| OTU-036 | Firmicutes | Bacilli | Lactobacillales | Enterococcaceae | *Enterococcus* |
| OTU-038 | Firmicutes | Clostridia | Oscillospirales | Ruminococcaceae | Unclassified |
| OTU-039 | Proteobacteria | Alphaproteobacteria | Rhodobacterales | Rhodobacteraceae | *Paracoccus* |
| OTU-040 | Proteobacteria | Alphaproteobacteria | Azospirillales | Azospirillaceae | *Skermanella* |
| OTU-041 | Proteobacteria | Alphaproteobacteria | Azospirillales | Azospirillaceae | *Skermanella* |
| OTU-042 | Desulfobacterota | Desulfovibrionia | Desulfovibrionales | Desulfovibrionaceae | *Desulfovibrio* |
| OTU-044 | Planctomycetota | Planctomycetes | Pirellulales | Pirellulaceae | *Blastopirellula* |
| OTU-045 | Proteobacteria | Gammaproteobacteria | Pseudomonadales | Cellvibrionaceae | *Cellvibrio* |

| | | | | | |
|---|---|---|---|---|---|
| OTU-046 | Firmicutes | Clostridia | Clostridiales | Clostridiaceae | *Clostridium* sensu stricto 1 |
| OTU-047 | Firmicutes | Bacilli | Bacillales | Bacillaceae | *Geobacillus* |
| OTU-048 | Firmicutes | Bacilli | Lactobacillales | Lactobacillaceae | *Lactobacillus* |
| OTU-049 | Firmicutes | Bacilli | Lactobacillales | Streptococcaceae | *Streptococcus* |
| OTU-051 | Bacteroidota | Bacteroidia | Flavobacteriales | Flavobacteriaceae | *Flavobacterium* |
| OTU-053 | Proteobacteria | Alphaproteobacteria | Sphingomonadales | Sphingomonadaceae | Unclassified |
| OTU-054 | Bacteroidota | Bacteroidia | Cytophagales | Hymenobacteraceae | *Rufibacter* |
| OTU-056 | Proteobacteria | Gammaproteobacteria | Enterobacterales | Colwelliaceae | *Colwellia* |
| OTU-058 | Actinobacteriota | Actinobacteria | Micrococcales | Dermacoccaceae | *Kytococcus* |
| OTU-059 | Firmicutes | Bacilli | Lactobacillales | Catellicoccaceae | *Catellicoccus* |
| OTU-060 | Proteobacteria | Alphaproteobacteria | Rhodobacterales | Rhodobacteraceae | *Sulfitobacter* |
| OTU-061 | Proteobacteria | Alphaproteobacteria | Rhizobiales | Beijerinckiaceae | *Methylobacterium-Methylorubrum* |
| OTU-062 | Firmicutes | Bacilli | Erysipelotrichales | Erysipelotrichaceae | Group ZOR0006 |
| OTU-063 | Firmicutes | Bacilli | Lactobacillales | Streptococcaceae | *Streptococcus* |
| OTU-066 | Proteobacteria | Alphaproteobacteria | Rhodobacterales | Rhodobacteraceae | Unclassified |
| OTU-070 | Actinobacteriota | Actinobacteria | Corynebacteriales | Corynebacteriaceae | *Corynebacterium* |
| OTU-071 | Actinobacteriota | Actinobacteria | Corynebacteriales | Corynebacteriaceae | *Corynebacterium* |
| OTU-072 | Proteobacteria | Gammaproteobacteria | Enterobacterales | Pseudoalteromonadaceae | *Pseudoalteromonas* |
| OTU-073 | Verrucomicrobiota | Lentisphaeria | Victivallales | Victivallaceae | Unclassified |
| OTU-074 | Proteobacteria | Gammaproteobacteria | Enterobacterales | Vibrionaceae | *Photobacterium* |
| OTU-075 | Firmicutes | Clostridia | Lachnospirales | Lachnospiraceae | *Blautia* |
| OTU-077 | Proteobacteria | Gammaproteobacteria | Enterobacterales | Pseudoalteromonadaceae | *Pseudoalteromonas* |
| OTU-078 | Bacteroidota | Bacteroidia | Flavobacteriales | Flavobacteriaceae | *Tenacibaculum* |

| | | | | | |
|---|---|---|---|---|---|
| OTU-079 | Bacteroidota | Bacteroidia | Cytophagales | Hymenobacteraceae | *Adhaeribacter* |
| OTU-080 | Proteobacteria | Alphaproteobacteria | Sphingomonadales | Sphingomonadaceae | *Erythrobacter* |
| OTU-081 | Firmicutes | Clostridia | Clostridiales | Clostridiaceae | *Clostridium* sensu stricto 1 |
| OTU-082 | Actinobacteriota | Actinobacteria | Corynebacteriales | Nocardiaceae | *Nocardia* |
| OTU-089 | Proteobacteria | Alphaproteobacteria | Defluviicoccales | Defluviicoccaceae | *Defluviicoccus* |
| OTU-095 | Actinobacteriota | Actinobacteria | Micrococcales | Micrococcaceae | *Rothia* |
| OTU-100 | Cyanobacteria | Cyanobacteriia | Cyanobacteriales | Chroococcidiopsaceae | *Chroococcidiopsis* |
| OTU-105 | Proteobacteria | Gammaproteobacteria | Salinisphaerales | Salinisphaeraceae | *Salinisphaera* |
| OTU-107 | Firmicutes | Clostridia | Peptostreptococcales-Tissierellales | Family_XI | *Anaerococcus* |
| OTU-111 | Firmicutes | Bacilli | Lactobacillales | Streptococcaceae | *Streptococcus* |
| OTU-112 | Firmicutes | Bacilli | Staphylococcales | Gemellaceae | *Gemella* |
| OTU-113 | Actinobacteriota | Actinobacteria | Corynebacteriales | Corynebacteriaceae | *Corynebacterium* |
| OTU-121 | Firmicutes | Bacilli | Mycoplasmatales | Mycoplasmataceae | *Mycoplasma* |
| OTU-122 | Patescibacteria | Parcubacteria | *Candidatus* Nomurabacteria | *Candidatus* Nomurabacteria | Unclassified |
| OTU-123 | Proteobacteria | Alphaproteobacteria | Rickettsiales | *Candidatus* Hepatincola | Unclassified |
| OTU-141 | Proteobacteria | Gammaproteobacteria | Enterobacterales | Vibrionaceae | *Vibrio* |

**Table S4**. PERMANOVA of the bacterial operational taxonomic units (OTU) richness in the midgut of 12 fish species from the Aegean Sea. Greece. Red indicates $p < 0.05$.

| | ***Diplodus annularis*** | ***Diplodus vulgaris*** | ***Euthynnus alleteratus*** | ***Mullus surmuletus*** | ***Pagrus pagrus*** | ***Pagellus erythrinus*** | ***Spondyliosoma cantharus*** | ***Scyliorhinus canicula*** | ***Sparisoma cretense*** | ***Scorpaena porcus*** | ***Scorpaena scrofa*** | ***Uranoscopus scaber*** |
|---|---|---|---|---|---|---|---|---|---|---|---|---|
| ***Diplodus annularis*** | | 0.366 | 0.063 | 0.889 | 0.293 | 0.176 | 0.200 | 0.113 | 0.829 | 0.369 | 0.249 | 0.252 |
| ***Diplodus vulgaris*** | | | 0.029 | 0.286 | 0.767 | 0.513 | 0.602 | 0.547 | 0.167 | 0.055 | 0.403 | 0.917 |
| ***Euthynnus alleteratus*** | | | | 0.208 | 0.030 | 0.029 | 0.030 | 0.029 | 0.029 | 0.059 | 0.027 | 0.098 |
| ***Mullus surmuletus*** | | | | | 0.175 | 0.114 | 0.145 | 0.145 | 0.620 | 0.486 | 0.121 | 0.304 |
| ***Pagrus pagrus*** | | | | | | 0.482 | 0.571 | 0.578 | 0.111 | 0.030 | 0.397 | 0.970 |
| ***Pagellus erythrinus*** | | | | | | | 0.947 | 0.971 | 0.087 | 0.030 | 0.913 | 0.630 |
| ***Spondyliosoma cantharus*** | | | | | | | | 0.913 | 0.090 | 0.029 | 0.972 | 0.602 |
| ***Scyliorhinus canicula*** | | | | | | | | | 0.084 | 0.029 | 1.000 | 0.628 |
| ***Sparisoma cretense*** | | | | | | | | | | 0.146 | 0.054 | 0.197 |
| ***Scorpaena porcus*** | | | | | | | | | | | 0.028 | 0.055 |
| ***Scorpaena scrofa*** | | | | | | | | | | | | 0.597 |
| ***Uranoscopus scaber*** | | | | | | | | | | | | |

**Table S5**. PERMANOVA of the bacterial operational taxonomic units (OTU) abundances in the midgut of 12 fish species from the Aegean Sea. Greece. Red characters indicate $p < 0.05$. Shaded cells indicate trophic habit: Brown: omnivore with a preference for animal material; Green: carnivore with a preference for fish and cephalopods; Blue: carnivore with a preference for decapods and fish; Purple: omnivore with a preference for plants.

| | *Diplodus annularis* | *Diplodus vulgaris* | *Euthynnus alletteratus* | *Mullus surmuletus* | *Pagrus pagrus* | *Pagellus erythrinus* | *Spondyliosoma cantharus* | *Scyliorhinus canicula* | *Sparisoma cretense* | *Scorpaena porcus* | *Scorpaena scrofa* | *Uranoscopus scaber* |
|---|---|---|---|---|---|---|---|---|---|---|---|---|
| *Diplodus annularis* | - | 0.085 | 0.114 | 0.026 | 0.031 | 0.029 | 0.059 | 0.200 | 0.030 | 0.056 | 0.114 | 0.5469 |
| *Diplodus vulgaris* | | - | 0.090 | 0.581 | 0.174 | 0.110 | 0.310 | 0.419 | 0.600 | 0.326 | 0.457 | 0.9156 |
| *Euthynnus alletteratus* | | | - | 0.057 | 0.030 | 0.057 | 0.459 | 0.408 | 0.028 | 0.027 | 0.264 | 0.3239 |
| *Mullus surmuletus* | | | | - | 0.207 | 0.143 | 0.087 | 0.057 | 0.147 | 0.060 | 0.170 | 0.369 |
| *Pagrus pagrus* | | | | | - | 0.403 | 0.028 | 0.118 | 0.111 | 0.090 | 0.112 | 0.2027 |
| *Pagellus erythrinus* | | | | | | - | 0.027 | 0.087 | 0.058 | 0.058 | 0.108 | 0.1946 |
| *Spondyliosoma cantharus* | | | | | | | - | 0.600 | 0.029 | 0.087 | 0.485 | 0.5063 |
| *Scyliorhinus canicula* | | | | | | | | - | 0.085 | 0.145 | 0.484 | 0.8893 |
| *Sparisoma cretense* | | | | | | | | | - | 0.058 | 0.091 | 0.3166 |
| *Scorpaena porcus* | | | | | | | | | | - | 0.144 | 0.2106 |
| *Scorpaena scrofa* | | | | | | | | | | | - | 0.6388 |
| *Uranoscopus scaber* | | | | | | | | | | | | - |

**Table S7**. Range and median values of the coefficient of variation of the most dominant OTUs and the number of shared between the individuals found in each fish species.

| | Coefficient of variation | | | |
|---|---|---|---|---|
| | Minimum | Median | Maximum | Shared OTUs |
| All 12 species | 126.1% | 496.4% | 666.0% | 61 (7.1%) |
| *Diplodus annularis* | 0.0% | 81.6% | 200.0% | 55 (12.6%) |
| *Diplodus vulgaris* | 0.0% | 116.1% | 200.0% | 38 (9.0%) |
| *Euthynnus alletteratus* | 16.3% | 93.6% | 200.0% | 36 (12.8%) |
| *Mullus surmuletus* | 10.3% | 113.5% | 200.0% | 29 (11.6%) |
| *Pagrus pagrus* | 0.0% | 50.8% | 173.2% | 45 (25.1%) |
| *Pagellus erythrinus* | 0.0% | 66.9% | 173.2% | 39 (21.0%) |
| *Spondyliosoma cantharus* | 8.1% | 68.8% | 200.0% | 34 (15.8%) |
| *Scyliorhinus canicula* | 8.4% | 100.9% | 200.0% | 45 (18.3%) |
| *Sparisoma cretense* | 15.2% | 91.3% | 200.0% | 27 (13.0%) |
| *Scorpaena porcus* | 10.5% | 115.5% | 200.0% | 42 (15.9%) |
| *Scorpaena scrofa* | 12.8% | 99.1% | 200.0% | 37 (15.4%) |
| *Uranoscopus scaber* | 24.7% | 117.4% | 173.2% | 34 (15.2%) |

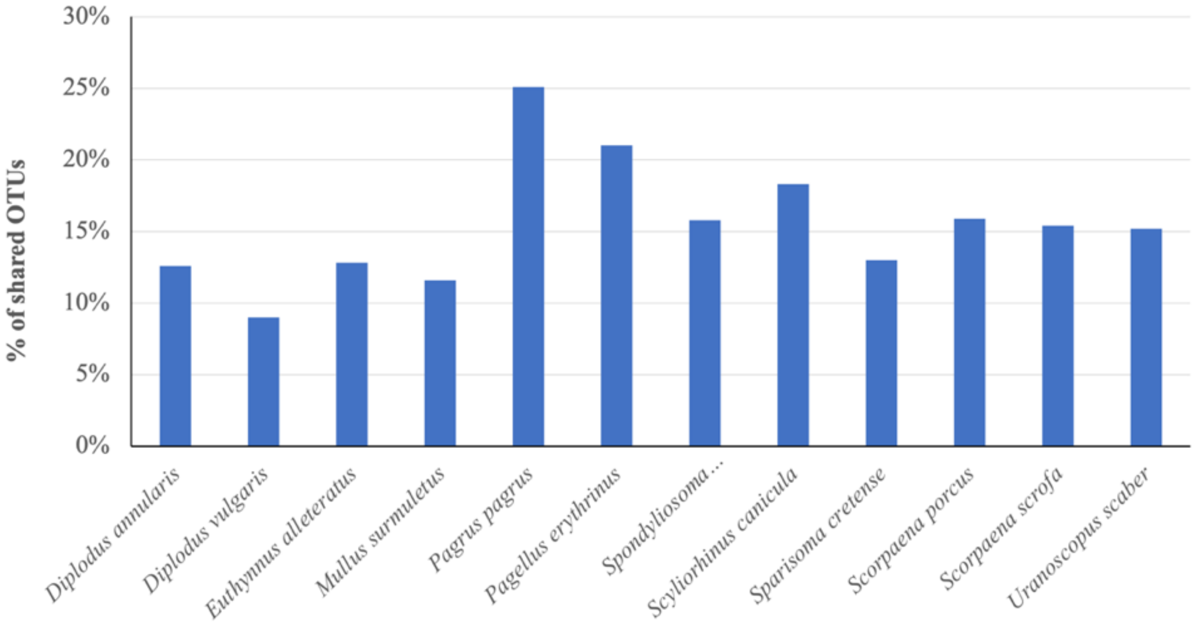


**Figure S1**. Shared operational taxonomic units in the midgut bacterial communities among the individuals of each of the 12 fish species from the Aegean Sea.

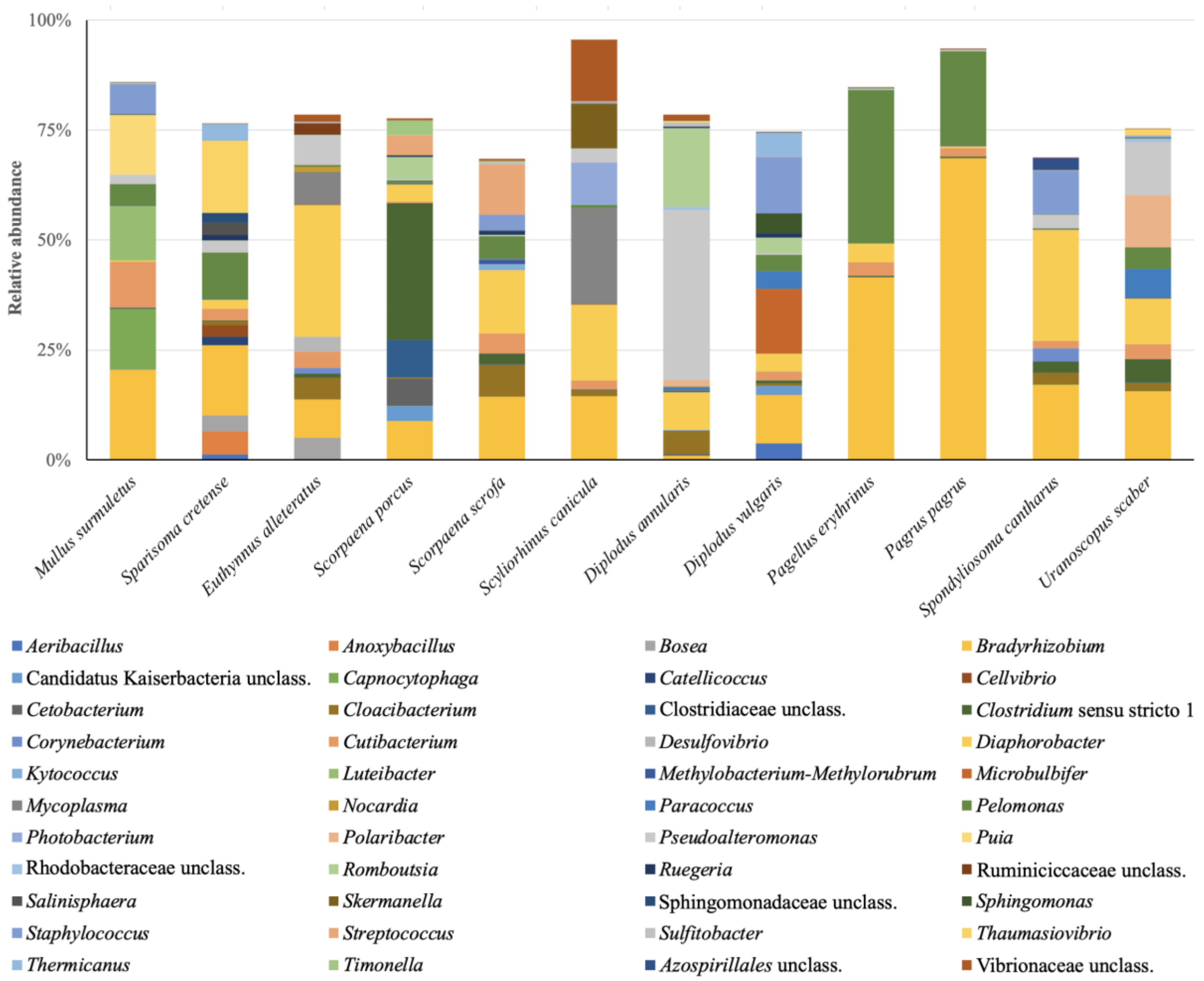


**Figure S2**. Taxonomic composition at the genus level or higher of the most dominant (≥70% cumulative relative abundance) bacterial operational taxonomic units of the 12 fish species. Note: multiple OTUs belonging to the same genus or higher taxon were pooled together.

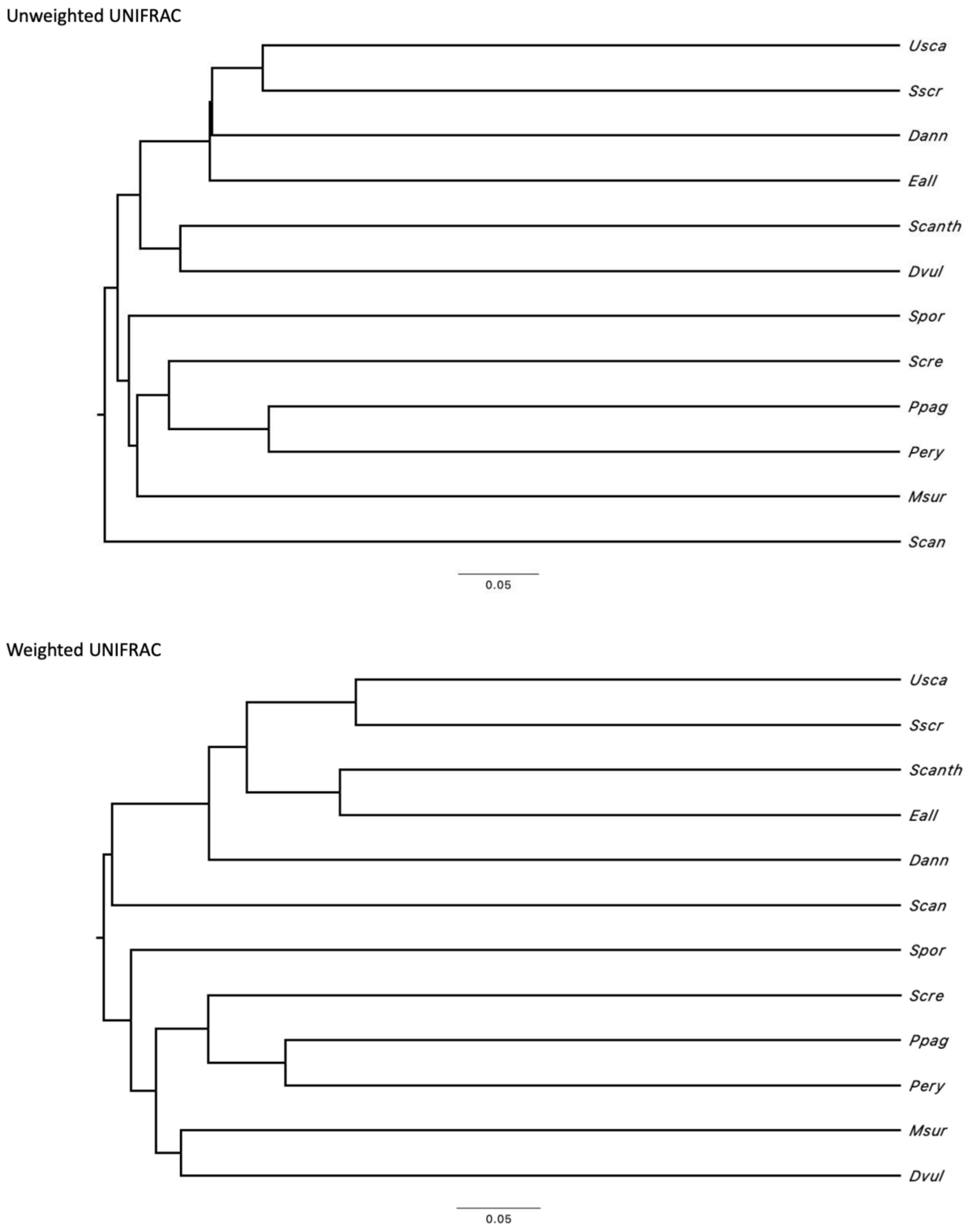


**Figure S3.** Unweighted and weighted UniFrac analysis of the 12 fish species' midgut bacterial microbiota.

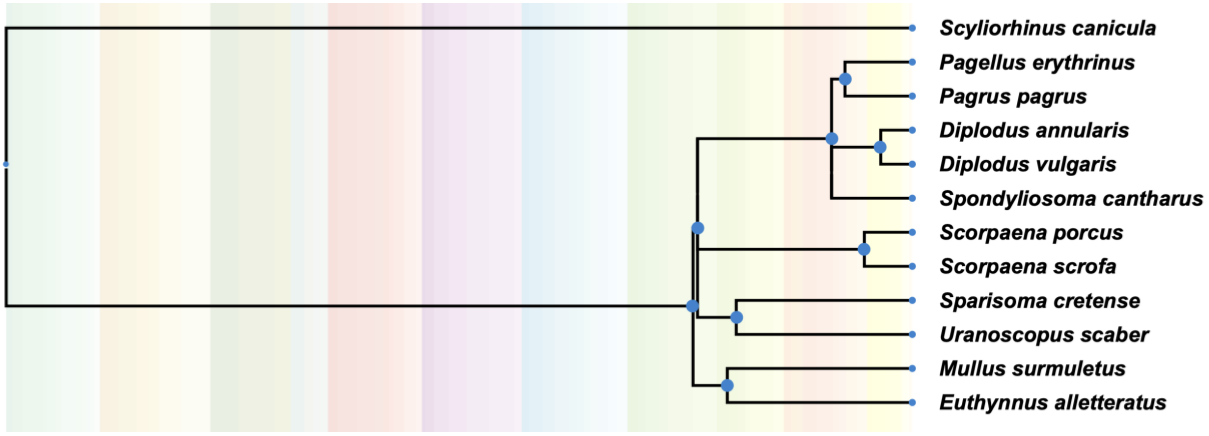


**Figure S4**. The phylogenetic relations of 11 of the 12 fish species from the Aegean Sea according to http://www.timetree.org. No adequate data exists for the phylogeny of *Uranoscopus scaber*.
